# Curvature Dependent constraints drive remodeling of epithelia

**DOI:** 10.1101/364208

**Authors:** Florian A. Maechler, Cédric Allier, Aurélien Roux, Caterina Tomba

**Affiliations:** Department of Biochemistry, University of Geneva, CH-1211, Geneva, Switzerland; Univ. Grenoble Alpes, CEA, LETI, DTBS, LISA, F-38000 Grenoble, France; NCCR Chemical Biology, University of Geneva, CH-1211, Geneva, Switzerland

## Abstract

Epithelial tissues are essential to keep a proper barrier for the organism. They usually have highly curved shapes, such as tubules or cysts. However, interplays between the environment and cell mechanical properties to set the shape are not known. In this study, we encapsulated two epithelial cell lineages, MDCK and J3B1A, into hollow alginate tubes and grew them under cylindrical confinement. Once formed, the epithelial MDCK layer detached from the alginate shell, while J3B1A layer remained adherent. Detachment resulted from contractile forces within cell layers that pulled cells away from the shell. We concluded that J3B1A cells have lower contractility than MDCK cells. As the pulling forces depend on the radius of the tube, we induced detachment of J3B1A cells by reducing the size of the hollow tube by two. Moreover, in bent tubes, detachment was more pronounced on the outer side of the turn, while extrusion occurred in the inner side, further highlighting the coupling between curvature and cell contractility.

## Introduction

In metazoan, epithelial tubules are structural features of organs to serve essential functions such as the transport of gases, liquids, metabolites or cells. Also, in organogenesis, epithelial tubules are often formed as intermediate templates for organogenesis. For example, gastrulation or neurulation are processes in which epithelia deforms into tubules to create the gut and the central nervous system respectively. The mechanism underlying the formation of organs and epithelial tubes formation, occurring during embryogenesis, are difficult to unravel because of the difficulty to entirely control (genetically, biochemically and mechanically) the microenvironment of these structures. Its morphogenesis relies on complex spatial rearrangement leading to complex structures (Keller, 2002; Salazar-Ciudad et al., 2003). Using tissue-engineering methods could considerably simplify the task by controlling at least the mechanical and the biochemical microenvironment of these tubes. This would help in deciphering what cellular properties dictate the final shape of the tissue grown in chemically and mechanically controlled microenvironment.

Even if in vitro studies of cell monolayers cultures on flat substrates have been broadly used, it often fails to recapitulate the three dimensions (3D) architecture of living tissues. The 3D architecture of epithelial cells allows for cells interactions with their microenvironment that are all lost when cultured in 2D or in diseases (Alford and Taylor-Papadimitriou, 1996; Broders-Bondon et al., 2018). In a 2D culture these interactions are poorly reproduced, which induces a loss of tissue-specific properties. A 2D culture can consequently neither support the tissue-specific functions of most cell types nor properly predict in vivo tissue functions (Greek and Menache, 2013).

To recapitulate a functional 3D organization, a simple method has been to culture specific cell types in hydrogels made of Extra-Cellular Matrix (ECM) components (Caliari et al., 2016). The interactions between cells and the ECM hydrogel create a complex network of mechanical and biochemical signals that are critical for normal cell physiology (Abbott, 2003; Griffith and Swartz, 2006; Pampaloni et al., 2007). However, the mechanical properties of such gels as well as their precise chemical composition are difficult to control or/and change (Beduer et al., 2015; Benenson and Lutolf, 2017). This has prompted the use of artificial hydrogels which composition and stiffness could be controlled accurately (Gjorevski et al., 2016). However, this method usually fails to apply geometrical or shape constraints on the growing tissue, as it is the case in vivo, where tissues grow under the pressure of surrounding organs. A growing effort is made for designing assays in which tissues are grown in microenvironments of controlled shape (Burute et al., 2017; Laurent et al., 2017). In accordance with this, the use of microfluidics in 3D cultures contributes to obtain controlled microenvironment where incorporate different cell types, respect the cell shape and mimic the mechanical and chemical diversity of living tissues with more accuracy (van Duinen et al., 2015).

Here, we present a cylindrical cell container made of alginate, in which cells can grow in a tube of similar dimensions than many in vivo tubular structures. The encapsulation technique used to produce these tubes already proved itself useful by producing hollow spheres to study the mechanics of tumour growth (Alessandri et al., 2013). In similar containers, however coated with Matrigel, a mix of ECM molecules, Neuronal Stem Cells can be differentiated into neurospheres, which are protected by the alginate shell, allowing for their manipulation (Alessandri et al., 2016). This technique controls many constraints that could impact epithelial morphogenesis and helps deciphering the specific impact of the microenvironment on cell growth, as well as tissue response to physical constraints (Roskelley et al., 1995).

With this cell container, we ought to understand how the cylindrical shape constraining the growth could affect the global organization and the final shape of two kinds of cell epithelial cell monolayers. We have selected two cell lines for their ability to form nicely organized epithelial layers, but with different cell size and appearance: Madin-Darby Canine Kidney (MDCK) and EpH4-J3B1A mammary gland epithelial cells. Both are some of the few cell lines that generate tubular structures in 3D cell cultures (Soulie et al., 2014). MDCK cells are a model cell-type in tissue mechanics and collective migration, which form monolayers with a relatively homogeneous cell aspect ratio. MDCK cells are able to form cysts, i.e. spherical and polarized monolayers with an inner lumen, from which tubulogenesis is induced when exposed for example to Hepatocyte Growth Factor (O’Brien, Zegers et al. 2002). J3B1A cells show slightly larger dimensions and have a more squamous cell aspect (Soulie et al., 2014). They usually form spheroidal cysts as well, but exhibit branching tubules in presence of low concentrations of Transforming Growth Factor-beta (Montesano et al., 2007).

## Results

### MDCK and J3B1A cells adapt their initial growth under tubular confinement

In this study, we confined and grew MDCK and J3B1A cell lines into a biocompatible and viscoelastic hollow tube made of alginate, a permeable (cut-off is 100 kDa) polymer with high potentials in biomaterials (Augst et al., 2006). By using 3D printed microfluidic chips, a co-axial three-layered jet flow was injected into a calcium bath (Fig. 1A). The microfluidic chip is a 3D printed device connecting three entry channels. These entries receive a flow from the connected syringe, respectively (i) a mix of cells, Matrigel and sorbitol (CS), (ii) sorbitol (IS) and (iii) alginate (AL). Using low-speed flow in the syringes allows the formation of droplets, at the exit point of the microfluidic device, falling in the calcium bath and producing alginate capsules (Alessandri 2016). However, when the fluxes of the syringes are appropriately increased and the nozzle is immerged in the calcium bath, the resulting continuous jet at the exit point of the chip produces a cylindrically constraint environment for the cells. A single tube can be elongated until the alginate stock is fully used, which corresponds to several meters of tubes. In this study, the length of the tubes filled in cells was of at least few centimeters in order to ensure an aspect ratio (length/radius) of about 100, avoiding eventual effects due to tube’s ends. Two different microfluidic devices were designed in this study, i.e. with exit canals of a diameter of 200 μm and 120 μm. For example, for tubes with J3B1A cells and 30% of Matrigel, these devices produced tubes an inner diameter of 428 ± 24 μm and 269 ± 13 μm respectively (mean ± SD as in the rest of the paper, otherwise noted). The diameter is smaller than the size at which diffusion of gazes and nutrients starts to be limiting (typically 500 μm).

**Figure 1.**
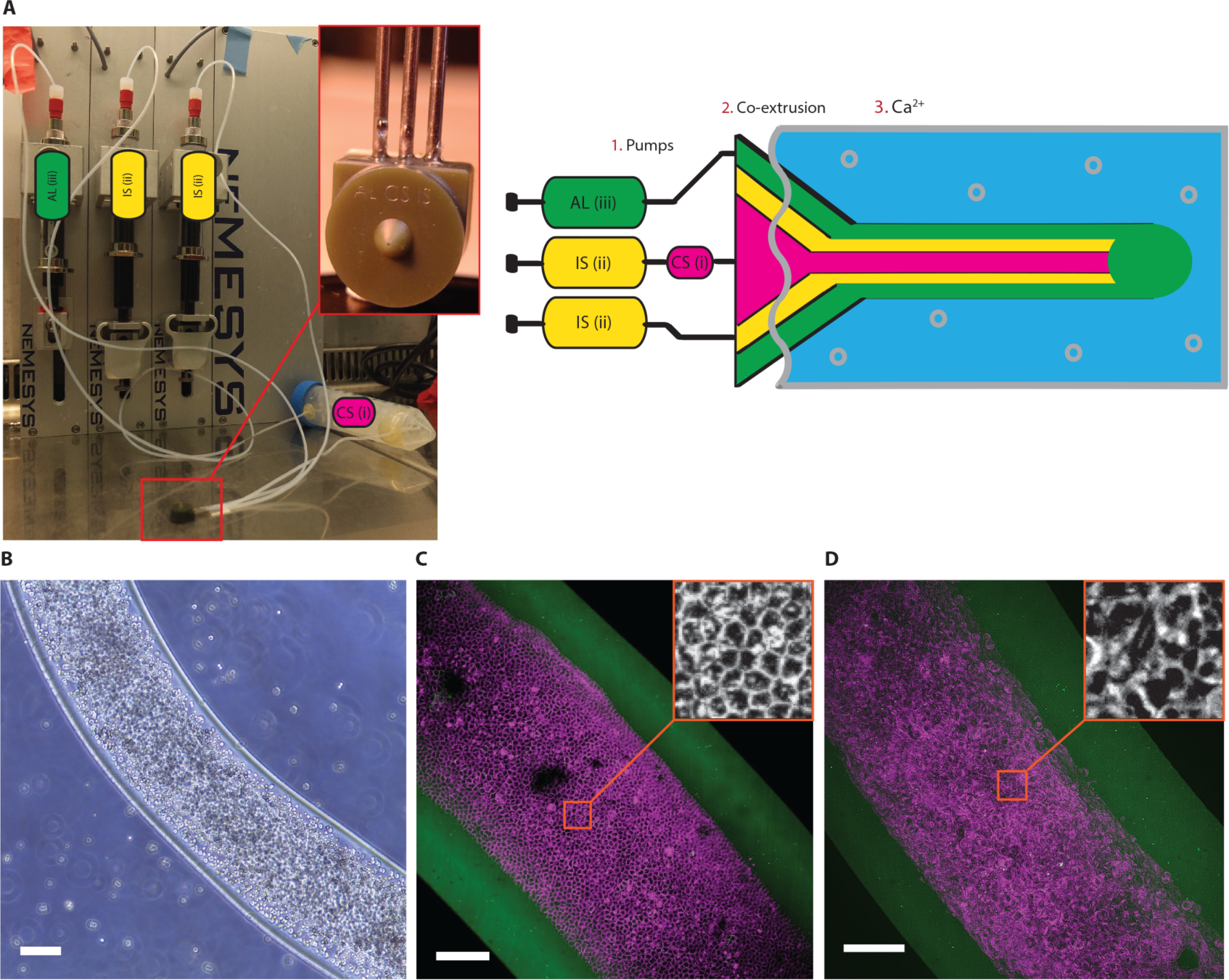
Set-up for the production of alginate tubes coated with a Matrigel layer on the inner side of the shell, where epithelial cells form a monolayer. (A) Picture (left) and diagram (right) of the set-up. 1. Pumps: set of the 3 syringes controlling the flow of the cell solution (CS (i)), the sorbitol (IS (ii)) and the alginate (AL (iii)). 2. Co-extrusion: 3D printed device (photograph in the inlet). 3. Ca^2+^: calcium gelation bath. (B) Brightfield image of a tube of MDCK cells at day 0, scale bar = 200 μm. (C-D) Maximum-intensity projections of 3D confocal images of a tube of (C) MDCK cells (20% Matrigel) at day 5 and of (D) J3B1A cells (40% Matrigel) at day 8. Alginate (g), cell membrane stained with CellMask (magenta), scale bars = 100 μm.

Since epithelial cells require efficient adhesion to their substrate to properly develop and since alginate does not provide a correct cell-adhesive interface (Lee and Mooney, 2012), a thin layer of Matrigel coated the inner surface of the hollow alginate tubes. This layer was prepared by adding Matrigel to the cold cell solution (4°C) and copolymerized with the alginate shell because of the temperature gradient (Alessandri et al., 2016). This leads to the enclosing of cells on tubular confinements, where cells initially flow freely inside the lumen of the tube. Then, they adhere to the Matrigel coating on the inner surface of the alginate wall (Fig. 1B). Through proliferation, both cell lines, MDCK (Fig. 1C) and J3B1A (Fig. 1D) cells, form clusters that merged into a monolayer, which closes upon reaching confluency. The tissue thus forms a cylindrical monolayer of cells with a lumen, and which dimensions are the ones imposed by the alginate shell. Differences between the tissue organizations between both cell lines can be underlined as the cellular arrangement allows MDCK cells to generate higher forces than J3B1A cells (Fig. 1C-D **inlets**). We first characterized the dynamics of proliferation of MDCK and JA3B1A cells into these alginate tubes.

### Three main stages characterize epithelia growth in alginate tubes

To follow epithelium formation within the alginate tubes at large as well as cellular scales, we took advantage of a lens-free microscope that allows the acquisition of large fields of view (up to 29.4 mm^2^, CMOS sensor) over long period of time (days) directly into the incubator (Allier et al., 2017). After an holographic reconstruction process, it is possible to obtain a phase image of the biological sample. The resolution is coarse (about 2-3 μm) and there is no sectioning ability but this was sufficient (Hervé et al., 2018) to follow growth of cells within alginate tubes. We could follow the evolution of the cell monolayer growing under cylindrical constraint over 5.5 days (Fig. 2A). First, after the monolayer has formed from merging clusters of cells, it spread onto the entire inner surface of the tube and created a lumen. Unexpectedly, upon reaching confluency, MDCK cell monolayer constricted, forming a narrower tube than the inner radius of the alginate shell, and causing the detachment of the monolayer from the alginate wall, probably disrupting its adhesion to the Matrigel layer.

**Figure 2.**
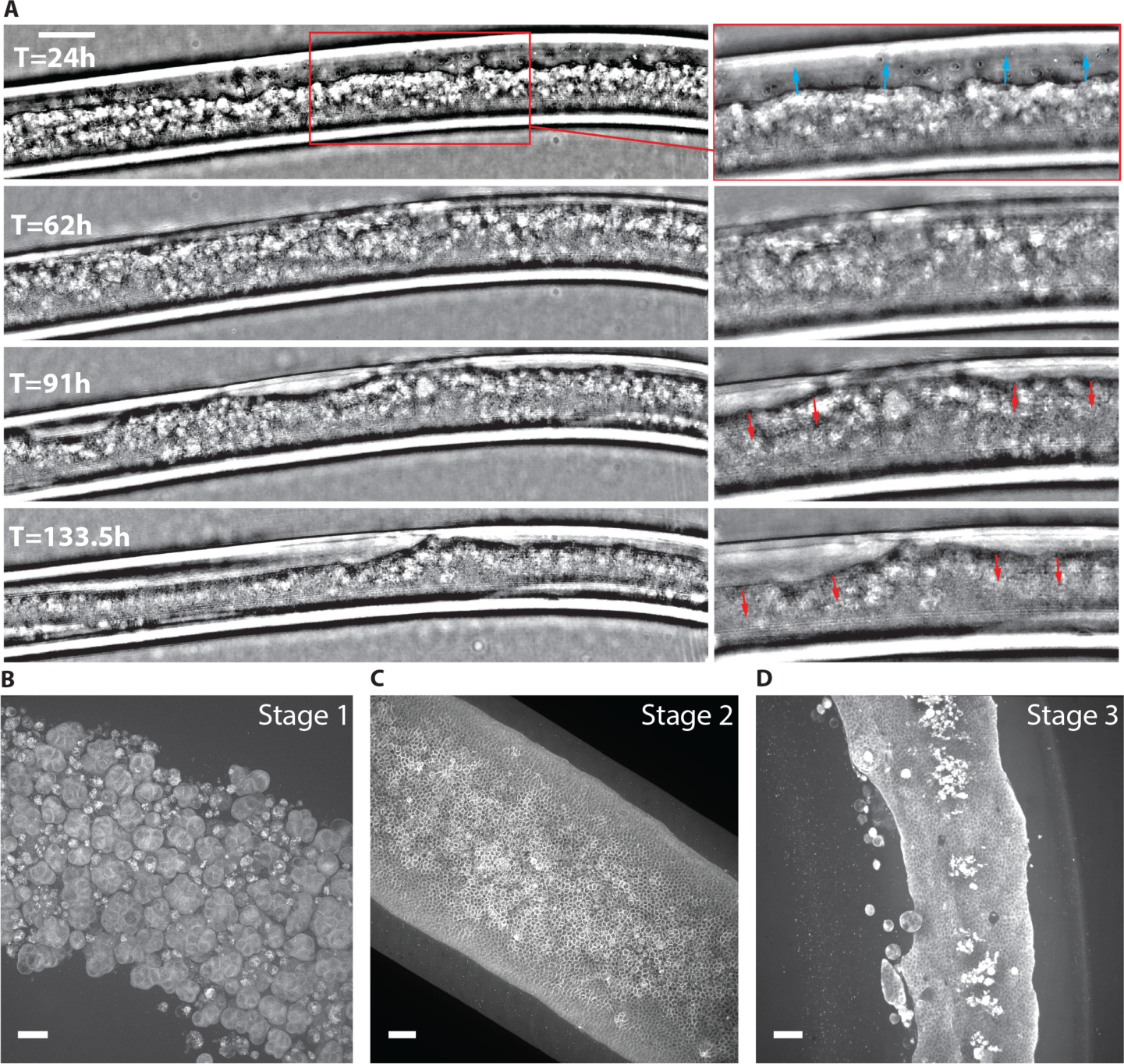
MDCK cell monolayer growth in alginate tubes. (A) Live imaging performed on a 5.5 days range with a lens-free microscopy (phase images). Frames in the inlets show monolyer progression (blue arrows, 24 hours after tube formation), tube closure at confluency (62 h) and detachment from the alginate wall (red arrows, 91 h – 133.5 h). Scale bar = 200 μm. (B-D): Maximum-intensity projections of 3D confocal images of the 3 stages of growth: (B) clusters (10% Matrigel, day 7), (C) monolayer (30% Matrigel, day 13), (D) shrunk monolayer (30% Matrigel, day 7). Cell membrane stained with CellMask (grey). Scale bars = 50 μm.

The growth evolution of the tissue can be described by three consecutive stages. Before a monolayer is formed, cells are arranged in clusters of 25-30 μm onto the inner surface of the tube wall (Fig. 2B, **stage 1**). Then, a proper monolayer forms while proliferating clusters of cells merge into a uniformly adherent cell monolayer onto the alginate wall (Fig. 2C, **stage 2**). Once at confluence, contraction of the monolayer occurs and lead to local or general loss of adhesion to the wall (Fig. 2D, **stage 3**). In order to investigate the possible effect of the Matrigel concentration on these stages, we first characterized the proportion of cells at each stage as function of time.

### MDCK cells present higher contractility compared to J3B1A cells

Cell shape can change through cell contractility, driven by the actomyosin cortex. Contractility strongly depends on ECM adhesion which transmits mechanical forces to the other cells of the tissue (Bielmeier et al., 2016). Thus, modulating the concentration of Matrigel may significantly influence tissue formation and behavior. We imaged alginate tubes containing MDCK and JA3B1 cells using bright field imaging for days after encapsulation, and for several Matrigel concentrations. Below 20% of Matrigel, MDCK cells cannot form a proper cell monolayer, and they mainly remain as clusters (Fig. 3A). At this concentration and below, the Matrigel does not coat homogeneously the entire inner surface of the alginate tubes, as observed by confocal imaging (**Supp.Fig. 3A-C**) and as previously published (Alessandri et al., 2016). However, J3B1A cells, even though with slower dynamics than for higher concentration of Matrigel, spread onto the entire surface of the alginate tubes with as low as 10% of Matrigel (Fig. 3B). But in this case, the cell monolayer rapidly detaches. This suggests that the holes present in the Matrigel coat below 20% concentration do not prevent JA3B1 cells from forming a monolayer and do not provide sufficient adhesion to the alginate substrate either. On the contrary, above 40% of Matrigel, J3B1A cells form spheroids, most probably because Matrigel is present in the lumen of the alginate tube at this concentration (**Supp.Fig. 3A-C**), while MDCK cells mostly stay as an adherent monolayer, but have an increased propensity to cell extrusion (see below).

**Figure 3.**
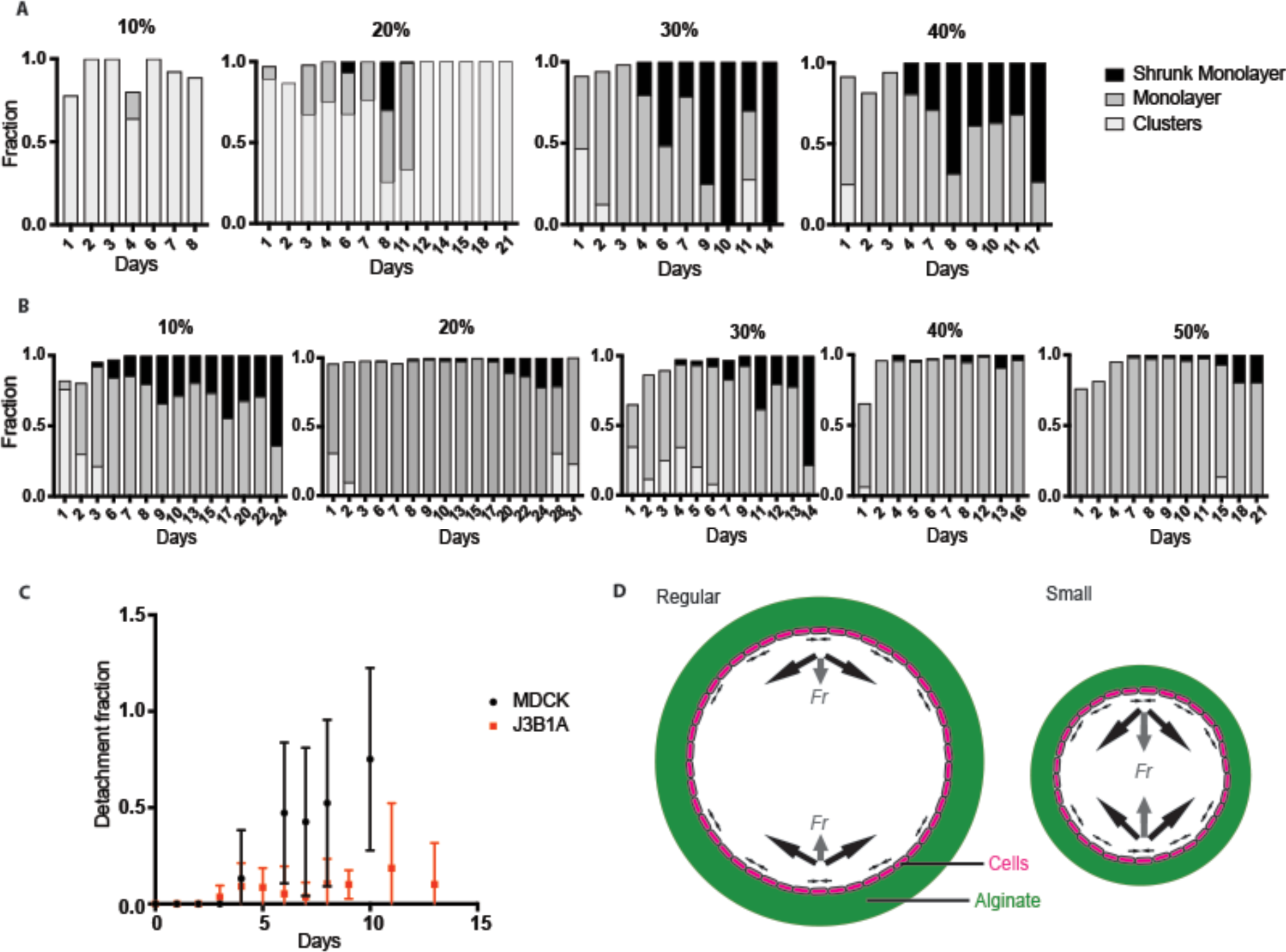
Comparison of the stages evolution for MDCK and J3B1A cells. (A-B) Histograms of the 3 stages of epithelial cells (A, MDCK; B, J3B1BA) in alginate tubes with different Matrigel concentrations (from 10 to 50%). (C) Averages of the detachment fractions for MDCK (black dots) and J3B1A (red squares) monolayers versus the days after tube formation. The fraction of detachment is calculated as the ratio between the length of detachment and the overall tissue length. Error bars = SD. (D) Diagram of regular (left) and small (right) tube orthogonal sections. Cells (magenta) forming a monolayer on the alginate wall (green) produce contractile forces (black arrows) with a bigger resulting force (“Fr”, grey arrows) in the small tubes.

For concentration between 20 and 40%, both MDCK and J3B1A cells form a monolayer within 3-4 days post-encapsulation, and this monolayer detaches rapidly for MDCK cells (4-6 days), while detachment is more limited for JA3B1 cells (Fig. 3A-B). For MDCK cells, the detachment fraction linearly increases over time, indicating a progressive contraction of the tissue once it has formed a complete monolayer (after 3 days). The detachment of the monolayer is therefore time dependent in MDCK cells, unlike J3B1A cells, which show a slight and almost constant detachment fraction after the monolayer formation (Fig. 3C). We propose that this may be explained by the stronger contractile forces that MDCK tissues can generate, as compared to J3B1A cells. In the smaller alginate tubes with J3B1A cells, we expected an increase of detachment compared to the regular ones at similar conditions as the inner curvature of the tube would result in higher force induced by the cell contractile forces (Fig. 3D).

In the tubular geometry, the contractile forces within the plane of the cell monolayer results in radial forces perpendicular to the tube surface and pulling the cells inward. One expects that the intensity of these radial forces is directly linked to the radius of the tube, the smaller the tube is, the larger the radial force is, if the contractile forces stay the same. Consistently with this reasoning, we observed both a faster appearance of shrunk J3B1A monolayers and an increase of the detachment fraction over time in the smaller alginate tubes (**Supp.Fig. 3D-E**).

Using the same reasoning, one expects that in curved tubes, detachment would be faster and larger in the outer side of the curve. We went on studying the detachment dynamics of cell monolayers in curved tubes.

### Monolayer detachment of MDCK cells is favored on the outer side of curved tubes

As tubes are formed and packed in petri dishes for cell culture, they adopt curved configurations. Long-term live-imaging of a tube of J3B1A cells in a turn of the tube showed the dynamics of the detachment process (Fig. 4A). 84 hours after the tube formation, shrinkage of the monolayer clearly appears at the outer part of the turn. This continues until the total detachment of the monolayer from the alginate wall on the outer side of the turn (130 hours). To quantitatively compare the frequency of detachment between the inner and outer side of a turn, we therefore took into account tubes with shrunk monolayers or cell extrusion from 4 days after formation, i.e. once cells reached confluency. By taking the ratio of the outer and inner detachment fraction (using 10^−5^ as 0 for no detachment of the inner side), we can distinguish three groups of values: under 0.01 (i.e. predominant detachment on the inner side of the turn), over 100 (i.e. predominant detachment on the outer side of the turn), and around 1 (similar detachment on both sides) (**Supp.Fig. 4A**). The asymmetry in monolayer detachment is the greatest at 30% Matrigel for MDCK cells, whereas for J3B1A cells, the peak occurs at 20%, but is loss completely at 50% of Matrigel (Fig. 4B). Surprisingly, when results from 20% to 40% Matrigel are combined, MDCK cells show a significant detachment in the outer side of turns as compared to straight tubes, whereas J3B1A cells do not show any difference between the straight and the bent tubes (Fig. 4C). But when the diameter of the tube is smaller, a clear difference is observed for J3B1A cells as well (**Supp.Fig. 4B-C**). This confirms that the pulling force resulting from the cell monolayer contraction is higher in smaller tubes, enhancing the curvature dependence of detachment.

While our results show that contractile forces are linked to curvature of the substrate to control detachment, the high variability of the tube curvature, and the fact that only short parts of the tube were straight, connecting two turns, made us look for a more controlled system of tube’s shape.

**Figure 4.**
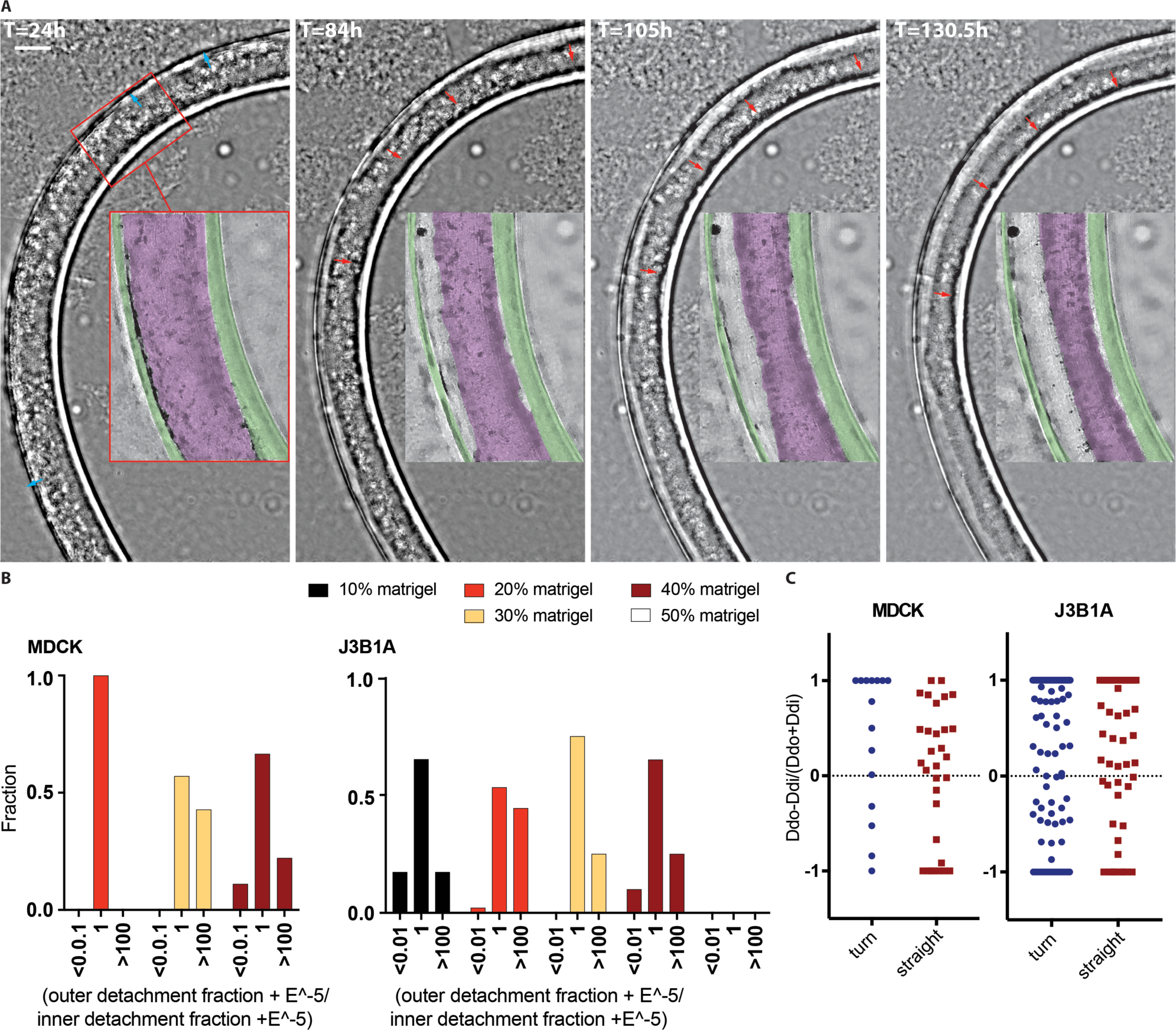
Epithelia detachment in curved tubes. (A) J3B1A cell monolayer growth in a curved alginate tube. Live imaging performed on a 5.5 days range with a lens-free microscopy (phase images). Frames in the inlets show monolyer progression (blue arrows, 24 hours after tube formation) and the progressive detachment from the outer part of the alginate wall (red arrows, 84 h – 130.5 h). Scale bar = 200 μm. (B) Histograms of the detachment fraction between the outer and the inner side of the bent tube of epithelia (MDCK, left; J3B1A, right) from 4 days after tube formation at different Matrigel concentrations (from 10 to 50%). (C) Fractions of detachment on the outer compared to the inner side of the tube turn for both cell lines. “Dd” = detachment fraction, “i” = inner side of the turn (blue), right side of the straigth tubes (red), “o” = outer side of the turn, left side of the stright tubes.

### Controlled curvature of the tube turn enhances the mechanical response also in J3B1A cells

Since bending tubes promote the monolayer detachment on the outer side of the turn, we evaluated quantitatively how curvature controls detachment by plotting the detachment fraction as a function of the curvature radius of the turns. We observed that for smaller radii, the difference of detachment between the outer and the inner side of the tube seemed to increase (Fig. 5A + **Supp.Fig. 5A**). However, the results were not statistically significant, and we looked for a way to obtain higher and more controlled values of curvature.

**Figure 5.**
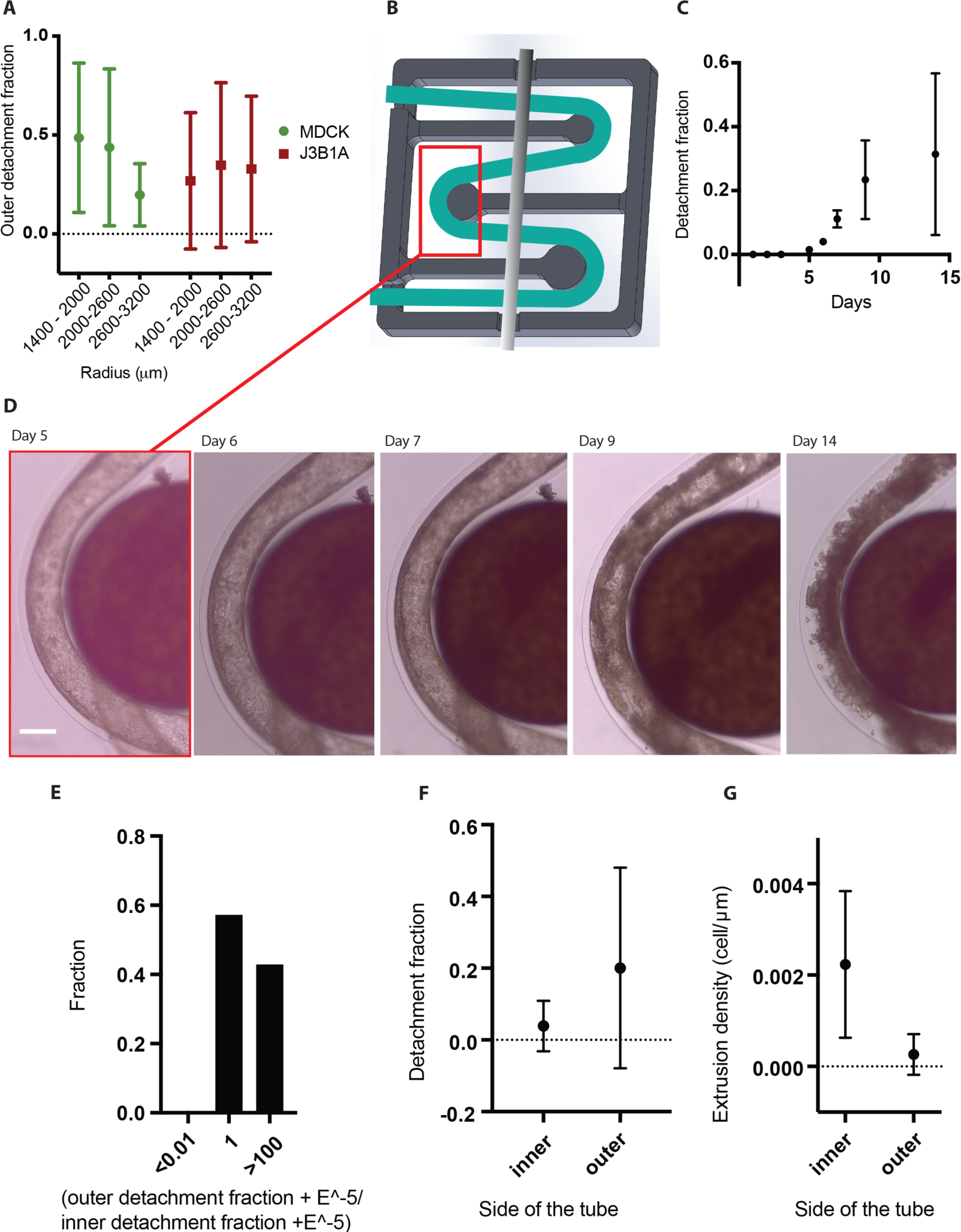
Epithelial monolayer growth in bent tubes. (A) Averages of the outer and inner detachment fraction of both cell lines (MDCK, green dots; J3B1A red squares) in bent tubes of different ranges of curvature radii of the turn. Error bars = SD. (B-G): J3B1A cells in small tubes under controlled curved constraint. (B) Design of the 3D printed tube holder (dark grey, 14 × 14 mm^2^), where the position of the alginate tube (green) is kept by a metallic bar (light grey). (C) Averages of the detachment fractions, on both sides of the tube, of J3B1A monolayers over time. Error bars = SD. (D) Bright field images of the evolution of the growth of J3B1A cells around a pillar of the tube holder. Scale bar = 200 μm. (E) Histogram of the detachment fraction of J3B1A monolayers from day 4. (F) Averages of the detachment fraction on the inner and the outer side of the turn from day 4. Error bars = SD. (G) Averages of cell extrusion densities on the inner and the outer side of the turn from day 4. Error bars = SD.

We designed a tube holder (Fig. 5B) in which tubes could be bent onto cylindrical pillars of a given radius, fixing the curvature of the turns. Using this device, we were able to obtain curvature radii as small as one millimeter. We then followed the tissue evolution of J3B1A cells in small tubes fixed in this controlled configuration of defined radius of curvature (Fig. 5C). This holder presents also the advantage to remove the need of using agarose to immobilize the tubes. With this holder, we observed a significant and a more regular increase of JA3B1A monolayer detachment over time (Fig. 5C-D), than on the previous results with small tubes (**Supp.Fig. 3E**), confirming that higher curvature can compensate the weak contractility of J3B1A cells to promote detachment. The asymmetry between the detachment fraction on both sides of the turn is also enhanced (Fig. 5E).

During these experiments, we consistently observed extrusion of cells towards the Matrigel layer in the inner side of the tube turns (Fig. 5F). In an epithelium, cell proliferation and cell removal must be balanced in order to keep its surface and shape constant. As local contraction of cells promotes extrusions of others in epithelium (Kocgozlu et al., 2016; Wu et al., 2015), we wondered if this curvature dependence of extrusion could have the same origin than detachment. Indeed, if extrusions are promoted by constriction of the epithelial cell actomyosin ring, extrusion should be enhanced by tissue curvature.

### Cell extrusion predominantly occurs in the inner side of the bending tubes

We systematically quantified the extrusion densities to fully characterize both cell lines behaviors in straight and bent tubes. Once at confluency, intercellular forces into the monolayer can create compressive patterns and extrude some cells, eventually forming clusters on its surface (Fig. 6A). Extrusion was greatly increased by higher concentrations of Matrigel in MDCK cells tubes (Fig. 6B), most probably because the Matrigel layer is thicker (**Supp.Fig. 3A-C**), reducing the constraints on extrusion of cells toward the alginate wall. It could also enhance contractility of cells through the stronger attachment of the monolayer to its ECM. MDCK cells show an average of extruded cell density, expressed as the number of extruded cells per micron, several times higher than J3B1A cells (Fig. 6B). Consistently with the higher detachment fraction of MDCK than of J3B1A tissues, this result reinforces the hypothesis of a higher contractility of MDCK than J3B1A cells.

**Figure 6.**
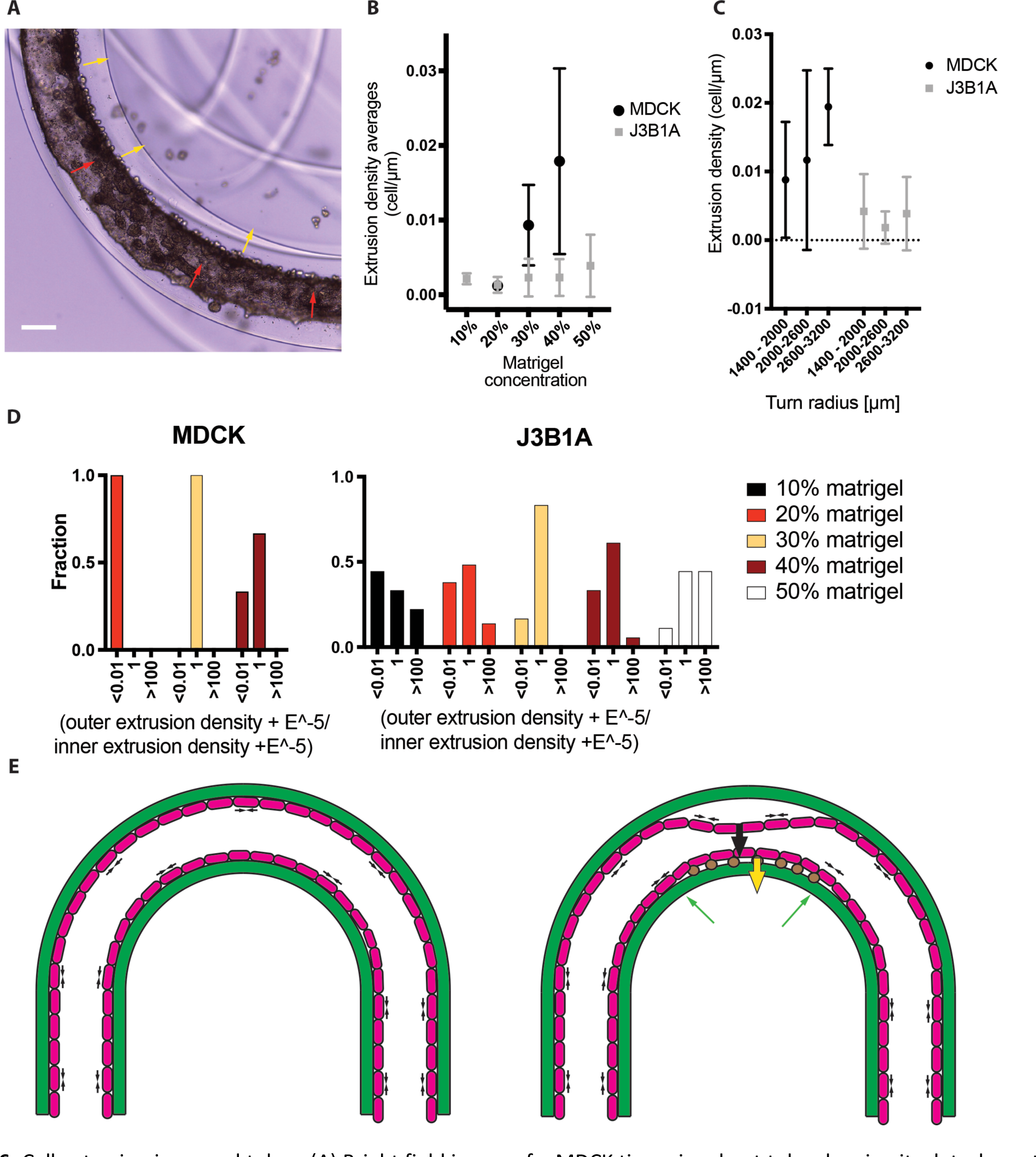
Cell extrusion in curved tubes. (A) Bright field image of a MDCK tissue in a bent tube showing its detachment on the outer side of the turn (red arrows) and the cell extrusions (yellow arrows) on the inner part. Scale bar = 200 μm. (B) Averages of the cell extrusion densities on the inner side of the tube turn for both cell lines (MDCK, black dot; J3B1A, grey squares) from day 4, at different Matrigel concentrations (from 10 to 50%). Error bars = SD. (C) Extrusion densities on the inner side of the turn used in (B) to compare different curvature radii of the tube turn. Error bars = SD. (D) Histograms of extrusion densities fractions between the outer and the inner side of the tube, for both cell lines (MDCK, left; J3B1A, right) after day 4 at different Matrigel concentrations (from 10 to 50%). (E) Diagram of the tube section showing the contractile lateral forces (black arrows) produced by the cell monolayer (magenta) formation in a curved alginate tube (green). On the right the resulting forces induce the monolayer detachment from the outer side of the turn (big black arrow) and cell extrusion (brown dots) on the inner side (yellow arrow), where the pulling force is balanced by the alginate wall (green arrows).

By selecting specifically regions in the turns of bent tubes, we observed that the extrusion density increases with decreasing radii of curvature of the tube turns (Fig. 6C). We also observed a striking asymmetry of cell extrusion upon confluency of MDCK cells (after a 100 hours) in curved tubes (Fig. 6A). Extrusion has the reverse probability than of the monolayer detachment (Fig. 4B). Indeed, we found for both cells lines, extrusion predominantly occurs in the inner side of the turns (Fig. 6D). However, this fact is reversed by higher concentration of Matrigel, as for 50%, J3B1A cells were mainly extruded on the outer side of the turns. The excess of adhesive molecules is probably at the origin of this behavior. In the smaller alginate tubes with J3B1A cells, the difference of extruded cell density between the inner and outer of a bent tube was significant. It confirmed the role that curvature has on the tissue cell reorganisation, as decreasing the tube’s radius allowed to increase both extrusion and extrusion asymmetry in the bent tubes (Fig. 6C, **Supp.Fig. 5B**).

These results, mirroring the ones obtained on monolayer detachment, support that the origin of extrusion is also cell contractility. The asymmetry observed on curved tubes (detachment of the monolayer on the outer side, extrusion on the inner side) may arise from the direction of the forces resulting from contraction: while the resulting force of contraction is pulling the monolayer off the alginate shell, leading to detachment on the outer side, the resulting force is pushing the cells towards the alginate shell, leading to cell extrusion in the inner side of the tube (Fig. 6E).

## Discussion

Here we show that epithelial cells growing under tubular confinement may result into cellular tissues of different size and shape because of the combination of cell properties and of the initial shape of the cell confiner. By growing cells into hollow tubes coated with Matrigel, we observed that cells form a complete monolayer after a few days. However, while the MDCK epithelium layer constricts soon after confluency and detaches from the tubular substrate to finally form a more constricted tube, the J3B1A cells mainly stay attached to the substrate, and adapt to the imposed shape. This is particularly well illustrated in bent tubes. In this situation, the constriction of MDCK cells leads to rapid detachment of cells from the substrate on the outer side of the tube turn, while it leads to extrusion of cells towards the Matrigel on the inner side of the turn. J3B1A cells monolayers do not display such drastic shape changes upon reaching confluency, probably because of their lower contractility. However, the asymmetric detachment and extrusion is recapitulated into thinner tubes (smaller diameter) or highly bent tubes of J3B1A cells. These results suggest that the forces for detachment or extrusion result from cell contractile forces summed over the curved surface, and that higher curvature can compensate for weaker contractile forces in order to provide the same detachment or extrusion forces.

By reducing the diameter of the tubes, we could make J3B1A cells to detach. Moreover, by bending the tubes, which corresponds to another curvature than the tube radius, we observed an asymmetry of cell detachment, MDCK cell layer detaching mainly from the outer side of turns. This asymmetry was mirrored by a significant proportion of cells extruded in the inner side of tubes’ turns. For JA3B1, asymmetry of detachment and extrusion was only observed in small tubes. All of our results are in agreement with the fact that curvature modulates the resulting force of contractile forces integrated over the surface of the entire epithelium. In fact, by changing curvature, we could modulate the intensity of the force that pulls the cells away from the alginate shell, or extrude them.

Matrigel concentration also affects the microenvironment and, because adhesion is essential for proper epithelial growth, the characteristics and development of the tissue. In both cell lines, Matrigel concentrations in the range of 20-40% induce increased detachment and extrusion asymmetries. At 10% Matrigel, there is no difference in detachment between both parts of the tube turn, and overall the detachment is limited. Indeed, if such Matrigel conditions confer a weaker attachment of the cell monolayer to its substrate, contractile forces are expected to be insufficient to detach the monolayer. On the reverse, at 50% Matrigel, the Matrigel layer is so thick that it does not provide the cells with a smooth surface to adhere on. This probably masks the effect of curvature, as the epithelium layer is not following the curvature of the tube.

Since these events result from self-imposed forces within the tissue, a characteristic property of each cell line, the final size/shape also depended on the cell type and on curvature. Indeed, when tubes are curved, the inner part of the turn has a higher extrusion density than straight tubes, and even higher density compared to the outer part of the turns (Fig. 6). The reverse asymmetry appears for the monolayer detachment, more frequent at the outer part of turns (Fig. 4).

Altogether, we show here that epithelial tissue can grow under cylindrical confinement, and that the final shape, as well as cell reorganization (monolayer detachment, extrusion) undergoing within the tissue depended on the adhesion to the confining substrate, shape and size of the substrate and cell type. Altogether, our findings provide instrumental information for the bioengineering field focusing on the control of tissue growth in vitro.

## Materials and methods

### 3D printing and assembling of the microfluidic device and of the tube holder

The microfluidic device was printed with a design of three co-axial channels as described in (Alessandri et al., 2016). A EnvisionTEC 3D printer was used in conventional working conditions. Prints were made with the HTM 140 V2 resin supplied by EnvisionTEC (Germany), which provides a resolution in the z axis of 25 μm. The process typically takes around 3 hours. The chip was extensively and gently rinsed with ethanol in order to rid of the liquid resin from the inner canals. It was then sonicated during 20 seconds, and exposed to UV light for 15 minutes. Three 19-gauge stainless steel segments of 2-3 cm with smoothed extremities were glued with epoxy resins (Loctite M-31CL, RS Components, Corby, UK) at the entry of each channel. PDMS was spread on the side of the hole of the chip tip to reduce surface tension. This treatment results to be crucial to get a regular jet at the exit of the device. Devices with holes of two different inner diameters have been printed, one of 200 μm (defined as “regular device”) and one of 120 μm (defined as “small device”). This last value resulted like the smallest diameter for which the 3D printer resolution allows to get functional channels.

The tube holder has a dimension of 14 mm × 14 mm in order to fit perfectly on the diameter of 20 mm of a glass bottom dish (MatTek, Bratislava, Slova Republic, Cat. # P50G-1.5-30-F). Two snicks are present on the lateral side in order to fit a metallic spindle which prevents the tube to float in the medium. The turns present in the maze have been designed at different radii of curvature (0.8 mm, 1 mm, 1.2 mm, 1.4 mm and 1.6 mm).

### Solutions and cell suspension preparation

Alginate (AL) was prepared by dissolving 2.5% wt/vol sodium alginate (Protanal LF200S; FMC) in water and by adding 0.5 mM of sodium dodecyl sulfate surfactant (SDS, ITW Reagents, Darmstadt, Germany, Cat. # A394). The solution was filtered with a 1 μm glass filter (Acrodisc®syringe filter, Pall Life Sciences, New York, U.S.A., Cat. # 4523). The alginate solution was labeled with AlexaFluor-488-TFP (Alexa-488-TFP, Invitrogen, Thermo Fisher Scientific, U.S.A, Waltham, Cat. # A30005) at 100[mg/L] and EDC-mediated coupled with sulfo-NHS. It was finally dialyzed with a Slide-A-Lyzer™ cassette 10K 12-20 mL capacity in water during a day, with the water changed every several hours. The intermediate solution (IS) was sorbitol (Merck, Darmstadt, U.S.A, Cat. # 56755) dissolved in milliQ water at a concentration of 300 mM and filtered by Steritop-GV with 0.22 μm pores (PVDF, Merck, Darmstadt, Germany, Cat. # MPSCGVT05RE). The cell solution (CS) at the innermost canal of the chip was prepared from a filtered suspension of cells (40 μm filter with a nylon cell strainer, Corning, Durham, U.S.A, Cat. # 431750) at a concentration of about 3 × 10^7^ cells/ml in a total amount of 300 μl of sorbitol and Matrigel (Corning, Durham, U.S.A, Cat. # 354234, Lot # 5173011). The solution was prepared according with the different Matrigel concentrations from 10 to 50 % vol/vol. For adhesive proteins (Matrigel) visualization, rhodamine labeled laminin (Red fluorescent, Cytoskeleton, Inc., Denver, U.S.A, Cat. # LMN01-A) was added to the Matrigel solution at a concentration of 10% vol/vol.

### Cell culture

EpH4-J3B1A mammary epithelial cells (J3B1A) and Madin Darby canine kidney cells (MDCK) were used. The cell growing medium for MDCK cells was Dulbecco’s Modified Eagle Medium (DMEM, Thermo Fisher Scientific, Waltham, U.S.A, Cat. # 61965026)supplemented with 10% fetal bovine serum (FBS, Thermo Fisher Scientific, Waltham, U.S.A, Cat. # 102701036), 1% of penicillin-streptomycin (PS, Thermo Fisher Scientific, Waltham, U.S.A, Cat. # 15140122) and 1% of Non-Essential Amino Acids Solution (NEAA, Thermo Fisher Scientific, Waltham, U.S.A, Cat. # 11140035). The J3B1A medium was Dulbecco’s Modified Eagle Medium (DMEM, Thermo Fisher Scientific, Waltham, U.S.A, Cat. # 21885025) supplemented with 10% HI Bovine serum (Thermo Fisher Scientific, Waltham, U.S.A, Cat. # 26170043), 1% of penicillin-streptomycin (PS, Thermo Fisher Scientific, Waltham, U.S.A, Cat. # 15140122).

Cell detachment from the culture flask has a different protocol for each cell line. For MDCK cells, trypsin-EDTA 0.05% (Thermo Fisher Scientific, Waltham, U.S.A, Cat. # 25300054) was incubated for 10 minutes at 37°C. For EpH4 J3B1A cells, Versene (Thermo Fisher Scientific, Waltham, U.S.A, Cat. #15040033) was incubated for 30 minutes at 37°C, and then trypsin-EDTA 0.05% (Thermo Fisher Scientific, Waltham, U.S.A, Cat. # 25300054) was added and incubated at 37°C for 5 minutes.

### Tube formation

Three pump-connected syringes (Analytical Science, Victoria, Australia, Cat. # 10MDR-LL-GT SGE) contained the solutions (alginate for the external channel and sorbitol for the intermediate and the inner channels) employed for the tube formation, and described above. The alginate flow was set at 115 [mL/h] with the standard device, and of 65 [mL/h] with the small device. The sorbitol flow was set at 105 [mL/h] with the standard device, and of 55 [mL/h] with the small device. The flow of the cell solution syringe was set at 90 [mL/h] with the standard device, and of 48 [mL/h] with the small device.

The alginate and the sorbitol solutions were loaded directly from the syringes to the connections on the device. The tube containing the cell solution was maintained in ice in order to keep the Matrigel under its gelation point (4°C). It was then plugged with the pump-connected syringe of sorbitol.

The gelation bath with calcium chloride 0.1 M (Sigma-Aldrich, St. Louis, U.S.A, Cat. # 449709) was pre-warmed at 37°C.

The tip of the device was dried before each tube production to avoid capillarity effects. Once the three pumps were simultaneously set off, the tip of the device was immersed in the calcium bath and a tube of about one meter could nominally be produced during the time when the cell solution was flowing out. Afterward, the calcium chloride was eliminated by aspiration with a sterile needle, washed with medium without serum, in order to avoid aggregates of precipitated proteins, and replaced by cell growing culture medium. The tubes were kept in an incubator with controlled environment (37°C, 5% CO2). Medium was changed every 2 to 3 days.

Between each experiment, the chip was extensively cleaned with deionized water, ethanol, deionized water, Phosphate-Buffered Saline (PBS, Thermo Fisher Scientific, Waltham, U.S.A, Cat. # 18912014), and deionized water again.

### Immunostaining

Tubes were fixed in 4% formaldehyde solution (Sigma-Aldrich, St. Louis, U.S.A, Cat. # F8775) in phenol red-free Minimum Essential Medium (MEM, Thermo Fisher Scientific, Waltham, U.S.A, Cat. # 51200046) for 20 minutes at room temperature (RT), washed twice with phenol red-free MEM Alpha Medium and stored at 4°C. After cell fixation, cells were stained with CellMask Deep Red Plasma Membrane Dye (Molecular Probes, Thermo Fisher Scientific, Waltham, U.S.A, Cat. # C10046) for 60 minutes at RT.

### Confocal imaging

Confocal images were obtained using Nikon Eclipse Ti microscope with a Zyla sCMOS camera with either an Apo LWD 40×/1.15 WI objective (Nikon) or a plan 20x/0.45 objective. An upright LSM 710 microscope with an Apochromat 20x/1.0 DIC objective (Carl Zeiss) was used to get confocal images of the empty tubes with green-fluorescent alginate and red-fluorescent laminin. Tubes were maintained in a glass bottom dish (MatTek, Bratislava, Slovak Republic, Cat. # P50G-1.5-30-F) with low melting temperature agarose (0.5 [g/L]) (Sigma-Aldrich, St. Louis, U.S.A, Cat. # A9414) covered by phenol red-free MEM (Thermo Fisher Scientific, Waltham, U.S.A, Cat. # 51200046).

### Live imaging

Live imaging was performed with a Cytonote lens-free video microscope (Iprasense). It is based on the lens-free imaging system described by Su et al. (Su et al., 2009) which was modified to perform continuous monitoring inside an incubator at a controlled temperature and humidity. It uses a CMOS image sensor with a pixel pitch of 1.67 μm and an imaging area of 6.4×4.6#x003D;29.4 mm^2^. Multiple wavelength illumination is provided by a multichip LEDs delivering red, green and blue illuminations. The RGB LEDs are located above a 150 μm pinhole, at a distance of approximately 5 cm from the cells. The phase image of the sample is obtained through a multispectral TV optimizer (Hervé et al., 2018). Live imaging was performed during 112 hours in an incubator with controlled environment (37°C, 5% CO2). Tubes were maintained in a glass bottom dish (MatTek, Bratislava, Slovak Republic, Cat. # P50G-1.5-30-F) with low melting temperature agarose (0.6 [g/L]) (Sigma-Aldrich, St. Louis, U.S.A, Cat. # A9414) covered by cell growing medium with phenol red-free MEM (Thermo Fisher Scientific, Waltham, U.S.A, Cat. # 51200046).

### Data collection and analysis

Values on the tissue states, i.e. monolayer detachment, clusters and cell extrusions, have been obtained from bright field images taken day after day throughout the evolution of the tissue growth. At least two images were taken for each tube, with a 5X PHO objective on a Leica S 40/0.45. Data from alginate tubes prepared under the same conditions (i.e. microfluidic device, Matrigel concentration and cell line) but at different days have been collected together. Image analysis was performed with ImageJ.

Fraction of detachment was calculated as the cautiously measured length of the alginate wall with an absence of monolayer due to its detachment (Dd) divided by the total length of the tube’s segment taken into account (Dt). To compare these detachment fractions between both side of the tube, we took the difference in length of the detached monolayer of the tube at the inner (Ddi) and the outer (Ddo) part of the bent tube divided by the total length of both the inner and outer segment (Dti and Dto).

